# Vitamin A and vitamin E metabolites comodulate amyloid-β aggregation

**DOI:** 10.1101/2021.10.30.466561

**Authors:** Priyanka Joshi, Sean Chia, Xiaoting Yang, Michele Perni, Johnny Habchi, Michele Vendruscolo

## Abstract

Alzheimer’s disease is characterized by the presence in the brain of amyloid plaques formed by the aberrant deposition of the amyloid-β peptide (Aβ). Since many vitamins are dysregulated in this disease, we explored whether these molecules participate in protein homeostasis by modulating Aβ aggregation. By screening 18 fat-soluble and water-soluble vitamins, we found that retinoic acid and α-tocopherol, two metabolites of vitamin A and vitamin E, respectively, affect Aβ aggregation both *in vitro* and in a *C. elegans* model of Alzheimer’s disease. We also show that effects of these two vitamin metabolites in combination can cancel each other out, suggesting that the complex composition of the cellular environment could have a protective role against protein aggregation through the simultaneous presence of aggregation promoters and inhibitors. Taken together, these results indicate that vitamins and their metabolites may be added to the list of components of the quality control system that regulate protein aggregation.

## Introduction

Alzheimer’s disease (AD) is the leading cause of dementia, a condition that affects over 50 million people worldwide and puts an enormous strain on our healthcare systems^1^. This disease is characterized by the accumulation of extracellular plaques formed of the amyloid-β peptide (Aβ) and of intracellular tangles formed by the tau protein^2–5^. Because of the toxicity associated with protein aggregation, living systems have evolved complex quality control mechanisms, collectively known as the protein homeostasis system, to control the presence of protein aggregates^4, 6, 7^. These mechanisms enable the regulation of protein synthesis, trafficking, interactions and degradation, and involve a wide range of cellular components, including enzymes^8^, molecular chaperones^7^, metabolites^9, 10^, and lipids^11^, which we are only beginning to understand in detail.

More generally, the protein homeostasis system is part of the wider cellular homeostasis system, which regulates all the cellular components, including lipids, carbohydrates, and metabolites. Increasing evidence suggests the presence of a close interplay between the lipid homeostasis and protein homeostasis systems, with different types of lipids having a wide range of effects on protein aggregation. One intriguing aspect of this interplay is the principle of ‘resilience in complexity’, according to which the opposite effects of different lipids on protein aggregation tend to cancel each other, so that lipid membranes of complex composition effectively buffer the aggregation-promoting roles of certain lipid types^12^.

In this work, we investigate whether a similar principle could be present for vitamins, with the aim of revealing a possible level of interplay between the metabolite homeostasis and the protein homeostasis systems. As a proof-of-principle, we focused on Aβ, as its abnormal aggregation is one of the underlying causes of Alzheimer’s disease^2–5^. We considered 18 fat-soluble and watersoluble vitamins (**Table 1**) as vitamin homeostasis has been reported to be dysregulated in aging, cognitive impairment and Alzheimer’s disease^13^. Further, we asked what are the microscopic mechanisms by which these metabolites act to modulate the aggregation of Aβ42. Finally, we show that the combination of an accelerator vitamin and an inhibitor vitamin cancels out their individual effects on the aggregation of Aβ42.

**Table 1.** List of the fat-soluble and water-soluble vitamins tested in the in vitro ThT-based screening assay. We tested 18 vitamins and found that vitamins A, D, E, K and B12 modulate Aβ42 aggregation.

| Vitamin | General name | Cellular location (HMDB) | Dysregulation status in AD (plasma/CSF level) | Effect on Aβ42 aggregation based on ThT binding kinetics |
| --- | --- | --- | --- | --- |
| Cobalamin | Vitamin B12 | membrane (predicted from logP) | decreased | Acceleration |
| Cholecalciferol | Vitamin D3 | cytoplasm, extracellular, membrane, mitochondria | decreased | Acceleration |
| Ergocalciferol | Vitamin D2 | cytoplasm, extracellular, membrane, mitochondria | decreased | Acceleration |
| Menaquinone | Vitamin K2 | not available | decreased | Inhibition |
| Menadione | Vitamin K3 | not available | decreased | Inhibition |
| Retinoic Acid | Vitamin A | cytoplasm, extracellular, membrane, nucleus, endoplasmic reticulum | decreased | Inhibition |
| α-Tocopherol | Vitamin E | cytoplasm, extracellular, membrane | decreased | acceleration |
| 4-Aminobenzoic acid | Vitamin B | not available | Not known | - |
| Biotin | Vitamin B7, Vitamin H | cytoplasm, extracellular, mitochondria, nucleus | decreased | - |
| Folic Acid | Folate, Vitamin M | extracellular | decreased | - |
| Niacinamide | Vitamin B3 | extracellular | decreased | - |
| D-Pantothenic acid hemicalcium salt | Vitamin B5 | extracellular, mitochondria | decreased | - |
| Pyridoxal hydrochloride | Vitamin B6 group | extracellular | decreased | - |
| Pyridoxamine dihydrochloride | Vitamin B6 group (biologically active form) | extracellular | decreased | - |
| Pyridoxine hydrochloride | Vitamin B6 group | extracellular | decreased | - |
| (-)-Riboflavin | Vitamin B2 | extracellular | not clear | - |
| Thiamine hydrochloride | Vitamin B1 | extracellular, membrane (predicted from logP), mitochondria | decreased | - |
| (+/-)-α-Lipoic acid | Vitamin-like antioxidant | Cytoplasm, extracellular, membrane (predicted from logP) | not clear | - |

Our results suggest that small molecule metabolites may assist proteins to remain in their soluble state, and that a homeostatic balance of metabolites and proteins underlies a robust molecular environment in a cell. Taken together, our study provides evidence that metabolite homeostasis acts to stabilize proteins against aggregation.

## Results

### An in vitro screen of 18 vitamins identifies modulators of Aβ42 aggregation

We carried out an *in vitro* aggregation screen based on thioflavin T (ThT), an amyloid-sensitive fluorescent dye (**Materials and Methods**). Our goal was to identify vitamins that modulate - either by accelerating or by inhibiting - the aggregation of Aβ42 (**Figure 1**). We considered 18 fat-soluble and water-soluble vitamins due to their reported dysregulation in Alzheimer’s disease^13^ (**Table 1**).

**Figure 1.**
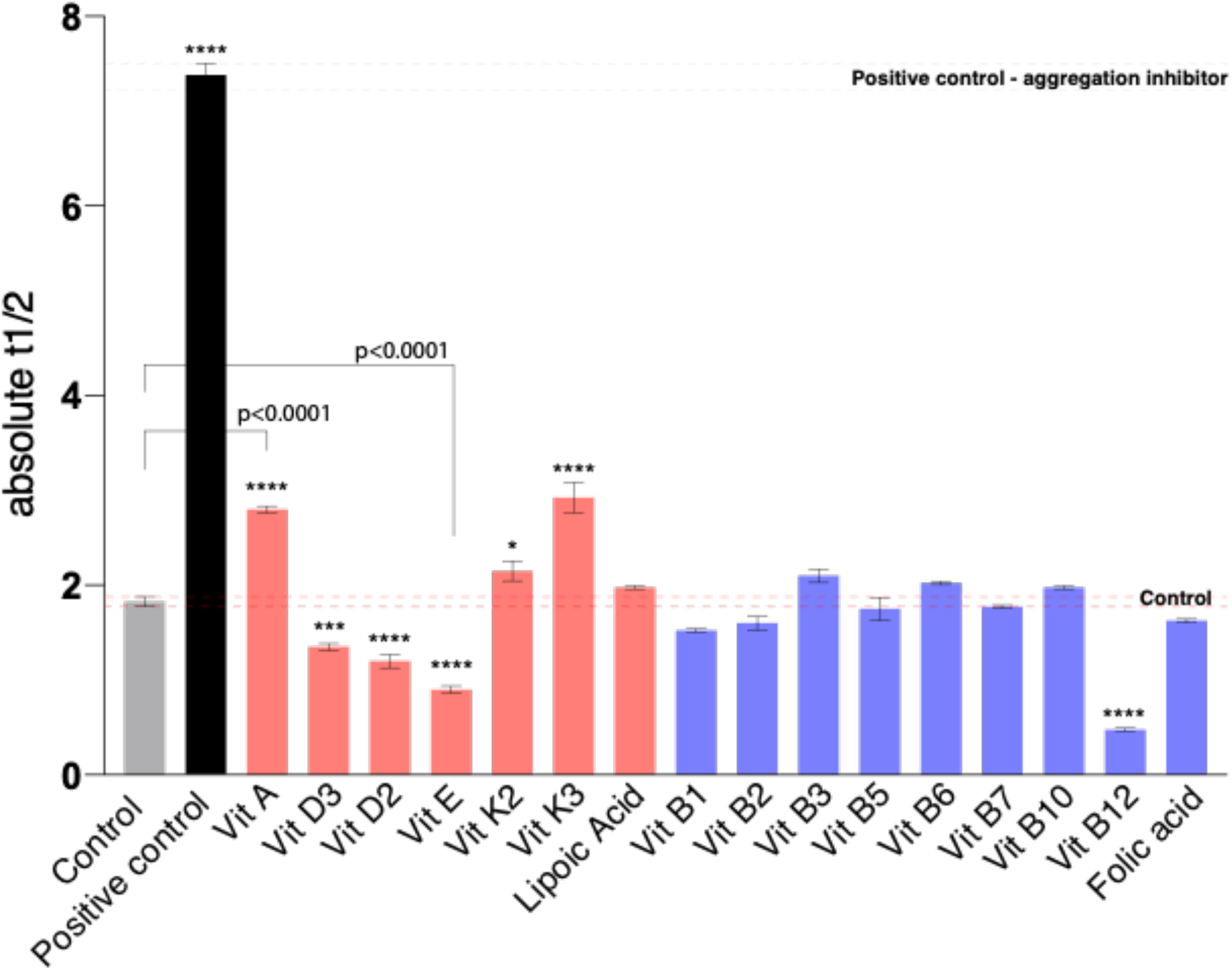
In vitro screening to identify vitamins that modulate Aβ42 aggregation. We tested 18 human endogenous fat-soluble (blue) and water-soluble (red) vitamins in a ThT-based screen to identify vitamins that either accelerate or delay the aggregation of Aβ42. By estimating the change in half-time of aggregation, we identified 7 vitamins as modulators of Aβ42 aggregation. The half-time is defined as the time at which the ThT signal has reached half of its final plateau value and is dependent on the initial monomer concentration. In the screen, the concentration of Aβ42 was 2 μM and that of vitamins was 20 μM. Plots are representative of 3 technical replicates, and we repeated the screening two times. Error bars depict the standard error of the mean (SEM). Statistics were performed using ordinary one-way ANOVA Dunnett’s multiple comparisons test (****, p-value<0.0001; ***, p-value=0.0005; *,p-value=0.03.

We found several vitamins that significantly altered the kinetics of Aβ42 aggregation in our screen at 20 μM metabolite concentration (10 molar equivalents, ME). At this concentration, these vitamins had significant effects on the half-time (t_1/2_) of Aβ42 aggregation, which is the time at which the ThT signal has reached half of its final plateau value (**Figure 1**). We found 3 vitamins (K2, K3 and A) that delayed Aβ42 aggregation, and 4 vitamins (B12, D2, D3 and E) that accelerated it (**Figure 1**). Although we found other vitamins in our screen with small significant effects on t_1/2_ (**Figure 1**), we focused on those with the greatest observed effects.

We next characterized each of these vitamins by individually carrying out the kinetic analysis at lower concentrations, which may be physiologically more relevant as intra- and extra-cellular concentrations of vitamins can be in the nanomolar to micromolar range. We found that at low concentrations, vitamins K2, K3, D2, D3 and B12 did not show significant effects (**Supplementary Figure S1**), while vitamins A (**Figure 2**) and E (**Figure 3**) induced reproducible and robust effects on Aβ42 aggregation.

**Figure 2.**
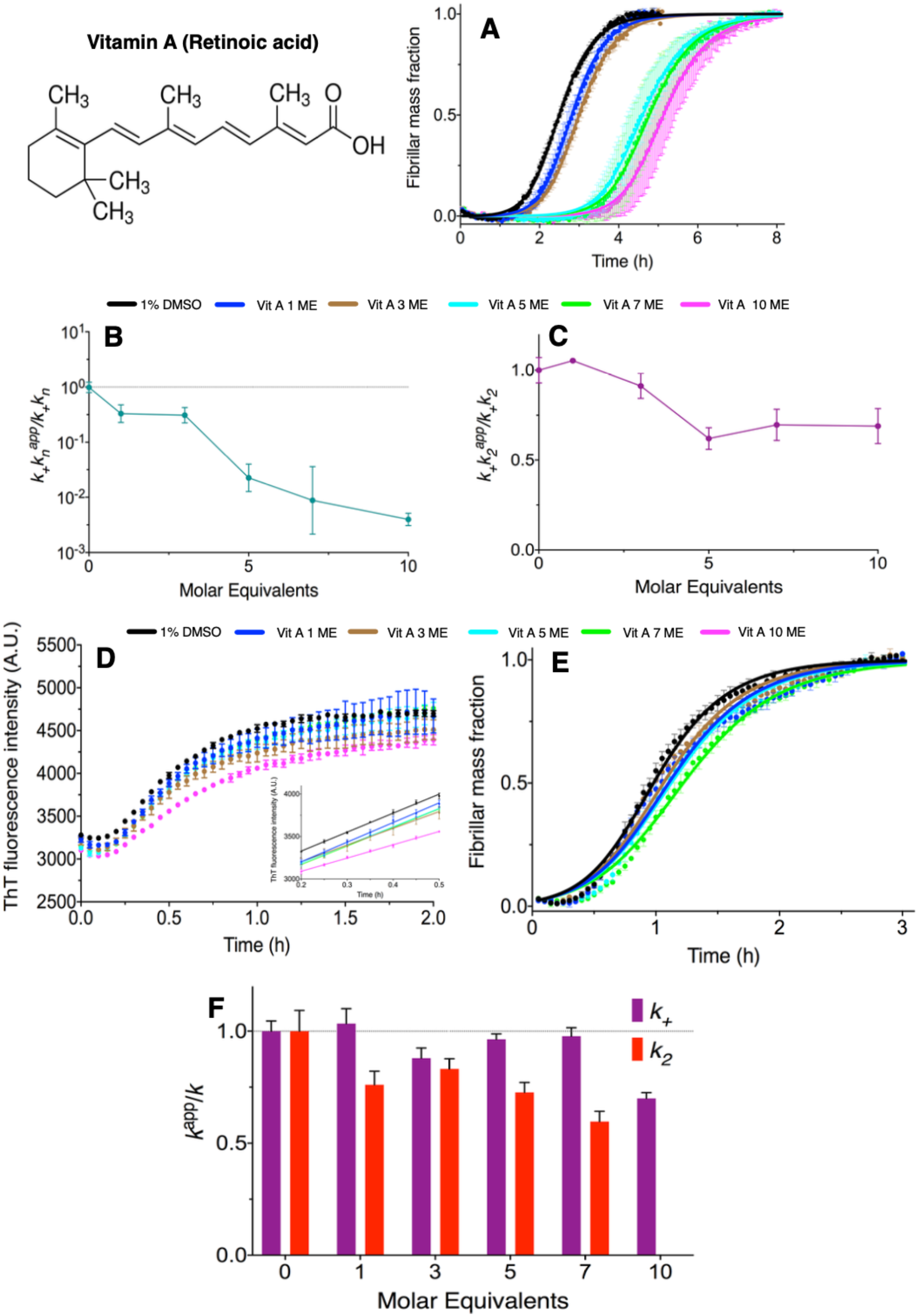
Vitamin A (retinoic acid) inhibits the aggregation of Aβ42 by affecting both primary and secondary pathways. **(A)** In an unseeded kinetic assay, we observed a dose-dependent inhibition of Aβ42 (in 1% DMSO) in the presence of vitamin A (1, 3, 5, 7 and 10 molar equivalents (ME), shown in different colours). **(B,C)** An analysis of the changes in the rate constants from the kinetic analyses in (**A**) with increasing concentrations of vitamin A shows a decrease in both primary (k_+_k_n_) and secondary (k_+_k_2_) pathways. **(D,F)** In a highly seeded kinetic assays, we saw that at 10 ME of vitamin A elongation is significantly affected by up to a 30% decrease. **(E,F)** We confirmed the decrease in k_2_ by performing the experiments in lightly seeded (2%) conditions at concentrations where elongation is not affected (1 to 7 ME). All fits were done using Amylofit^17^. Plots are representative of 3 technical replicates, and we performed this assay two times.

**Figure 3.**
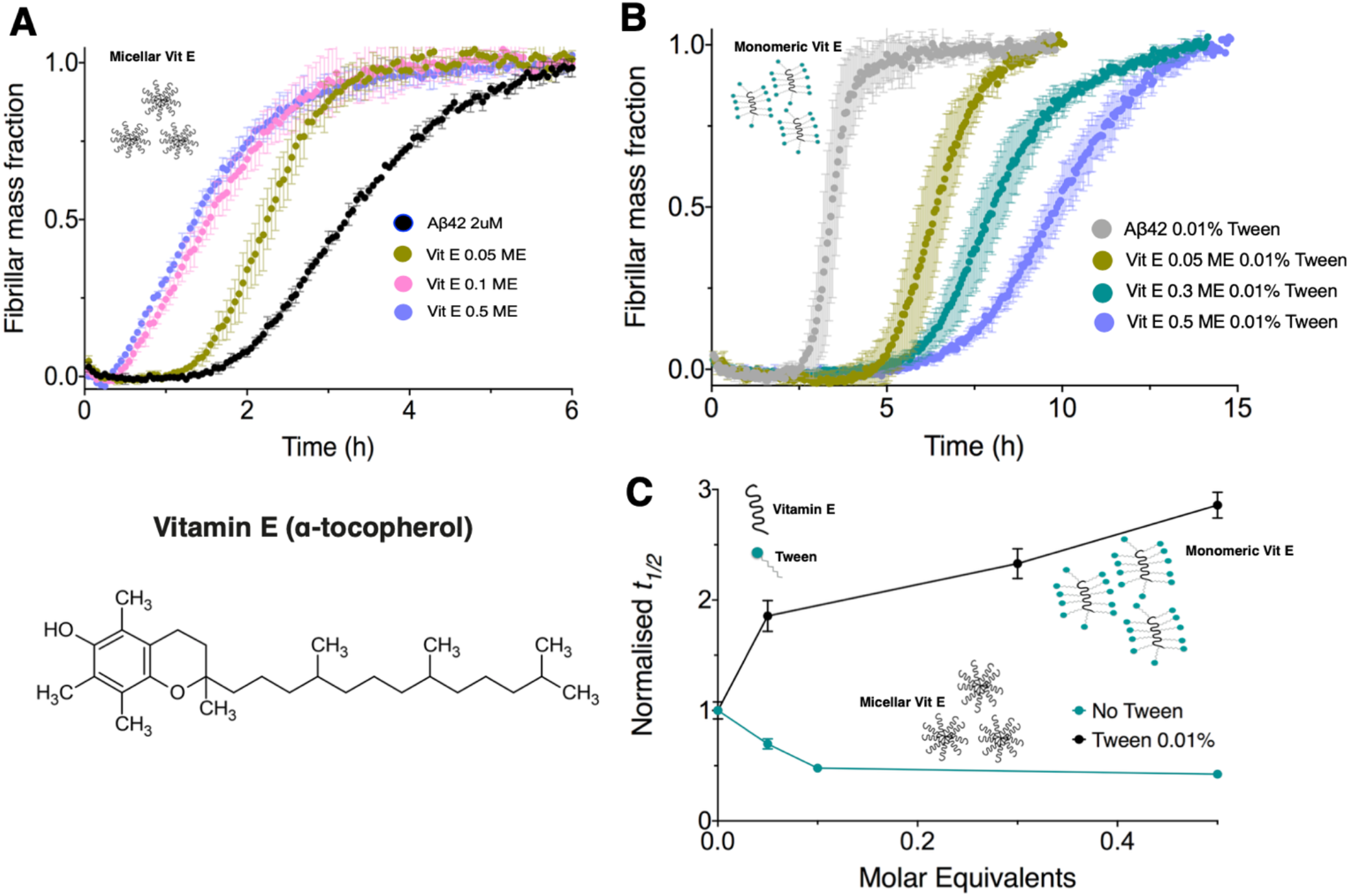
Vitamin E in a micellar form accelerates Aβ42 aggregation and inhibits it in a monomeric state. **(A,C)** We observed that vitamin E speeds up Aβ42 aggregation in a concentration-dependent manner when in a micellar form. **(B,C)** Vitamin E in a monomeric form (in the presence of Tween 0.01%) inhibits the aggregation of Aβ42 in a concentration-dependent manner. Plots are representative of 3 technical replicates, and we performed this assay two times.

### Vitamin A inhibits Aβ42 aggregation by affecting both primary and secondary pathways

The microscopic processes that govern the aggregation of Aβ42 are^4, 14–16^: (i) primary nucleation, which depends on the concentration of free monomers and proceeds with a rate constant k_n_, (ii) secondary nucleation, which depends both on the concentration of free monomers and on the concentration of aggregate mass and proceeds with a rate constant of k_2_, and (iii) elongation, where free monomers add to the growth-competent ends of existing fibrils and proceeds with a rate constant k_+_. We observed a dose-dependent inhibition of Aβ42 in the presence of vitamin A (retinoic acid) (unseeded kinetics, **Figure 2B**). Fitting the data using Amylofit^17^ shows that there is a decrease in k_+_k_n_ and a decrease in k_+_k_2_ (**Figure 2B,C**). We next decoupled the effects of k_+_ and k_n_ in the combined rate constants. Using highly seeded kinetics, we sought to use the growth rate at early timescales (**Figure 2D**, inset) to reveal effects on the elongation rate constant^14^, and we found that the elongation rate constant was only significantly affected at 10 ME of vitamin A, where a 30% decrease in k_+_ was observed (**Figure 2D,F**). Thus, for lower ME of vitamin A (i.e., 1 to 7 ME), a dose-dependent inhibition in primary pathways of Aβ42 as observed in the unseeded kinetics is likely due to a significant decrease in k_n_ (**Figure 2B**).

Next, we confirmed the decrease in k_2_ by performing the experiments in lightly seeded conditions (2% seeds), at molar equivalents where elongation is not significantly affected (1 to 7 ME, **Figure 2E**). We observed that the kinetics is also inhibited under these conditions (**Figure 2E, F**). Note that we did not fit the 10 ME results in these experiments, as vitamin A affects elongation at this high concentration, and its challenging to decouple the effects of elongation vs. secondary nucleation. Taken together, our data suggest that vitamin A (retinoic acid) inhibits the aggregation of Aβ42 predominantly by a combined effect on the primary and secondary processes.

### Vitamin E has opposite effects on Aβ42 aggregation in its monomeric and micellar forms

We next quantified the effects of vitamin E on Aβ42 aggregation. We observed that when vitamin E was added in DMSO alone it sped up the aggregation of Aβ42 (**Figure 3A,C**). Using dynamic light scattering (DLS), we observed that vitamin E forms micelles (**Figure 4**) in such conditions, a process that can provide a surface promoting the aggregation of Aβ42. However, when vitamin E is added in the presence of Tween 20 (0.01%), where it is solubilized, we observed that it inhibits the aggregation (**Figure 3B,C**). For comparison, when we added Tween 20 (0.01%) to the stock solution of vitamin A to break down potential higher order metabolite-assemblies, we did not find any differences in the inhibition of Aβ42 (**Figure S2**). DLS data confirmed the breakdown of micelles formed by vitamin E on addition of 0.01% Tween 20 (**Figure 4**). Indeed, a combination of accelerating and inhibitory properties of vitamin E depending on its micellar or monomeric states respectively is observed. In this context, we did not attempt to understand the effects of vitamin E on the individual microscopic steps of Aβ42 aggregation.

**Figure 4.**
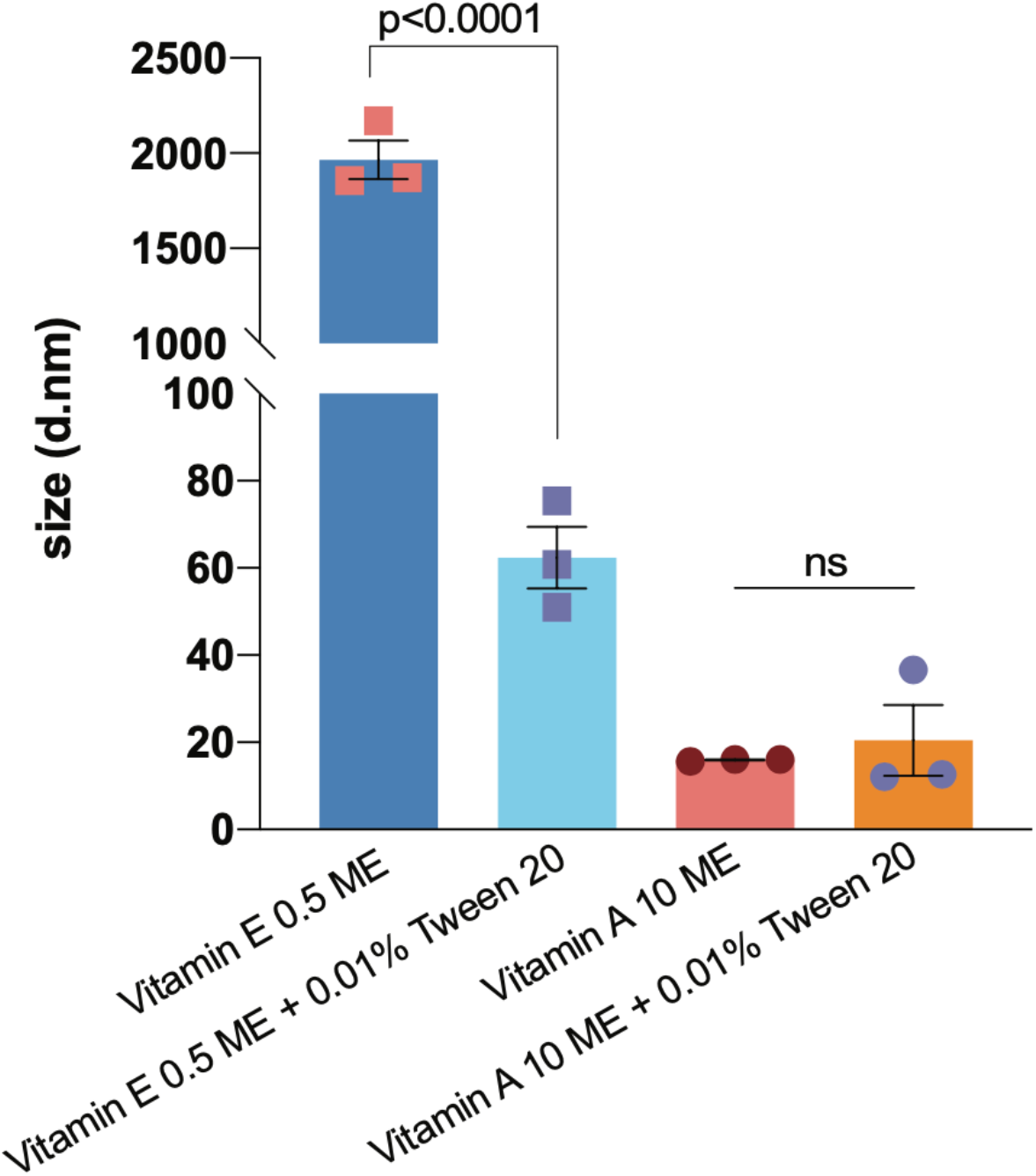
Dynamic light scattering (DLS) shows that vitamin E at 1 μM (0.5 ME) forms micelles which can be broken down by the addition of 0.01% Tween 20. By contrast, the size of vitamin A in solution is not affected by the addition of Tween 20. Statistical comparisons are shown using an unpaired t-test, p-value<0.0001. Data shows three independent replicates.

### Combining vitamin A and vitamin E leads to a zero net effect on Aβ42 aggregation

We next carried out the Aβ42 ThT-based aggregation assays in the presence of a mixture of retinoic acid and α-tocopherol. We thus observed that the two metabolites cancelled out their individual effects (**Figure 5**). Although α-tocopherol in micellar form at high concentrations accelerates Aβ42 aggregation (**Figure 4**), we found that the strong inhibitory role of retinoic acid subdued this effect even at some of the high α-tocopherol concentrations. After testing for a combination of concentrations in a mixture, we found a concentration range (0.15-0.25 ME α-tocopherol and 0-7 ME retinoic acid) within which the net effect of retinoic acid and α-tocopherol was zero on the aggregation profile of Aβ42 (**Figure 5)**.

**Figure 5.**
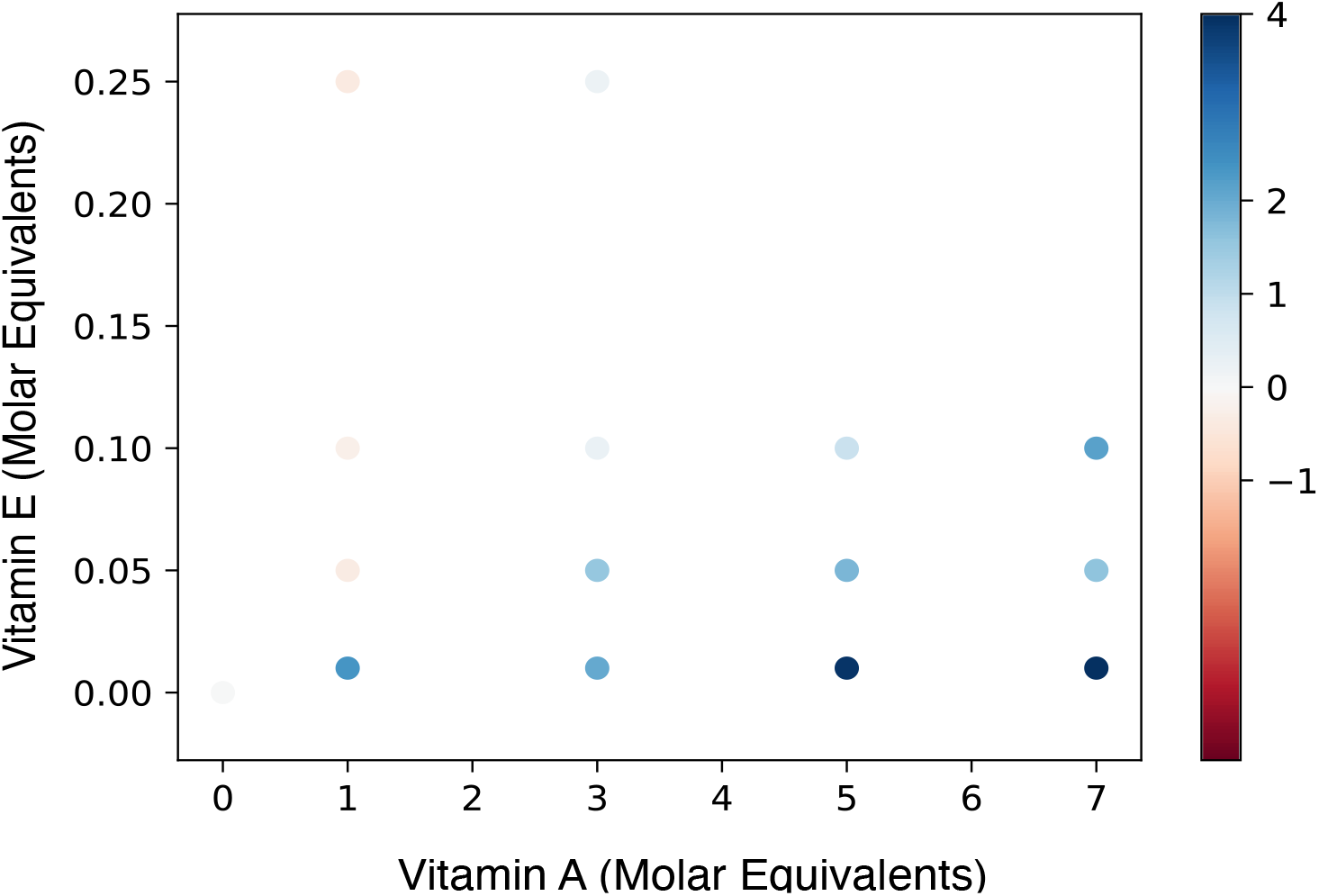
Heat map of the half-time (t_1/2_) of Aβ42 aggregation of different mixtures of vitamin A and vitamin E. The x-axis shows the molar equivalents of vitamin A added to the molar equivalents of vitamin E on the y-axis. The map shows the range of concentrations at which the individual effect of each metabolite is cancelled out by the other, thus having no net effect (zero-sum; equal to all data points representing 0) on Aβ42 aggregation.

**Figure 5.**
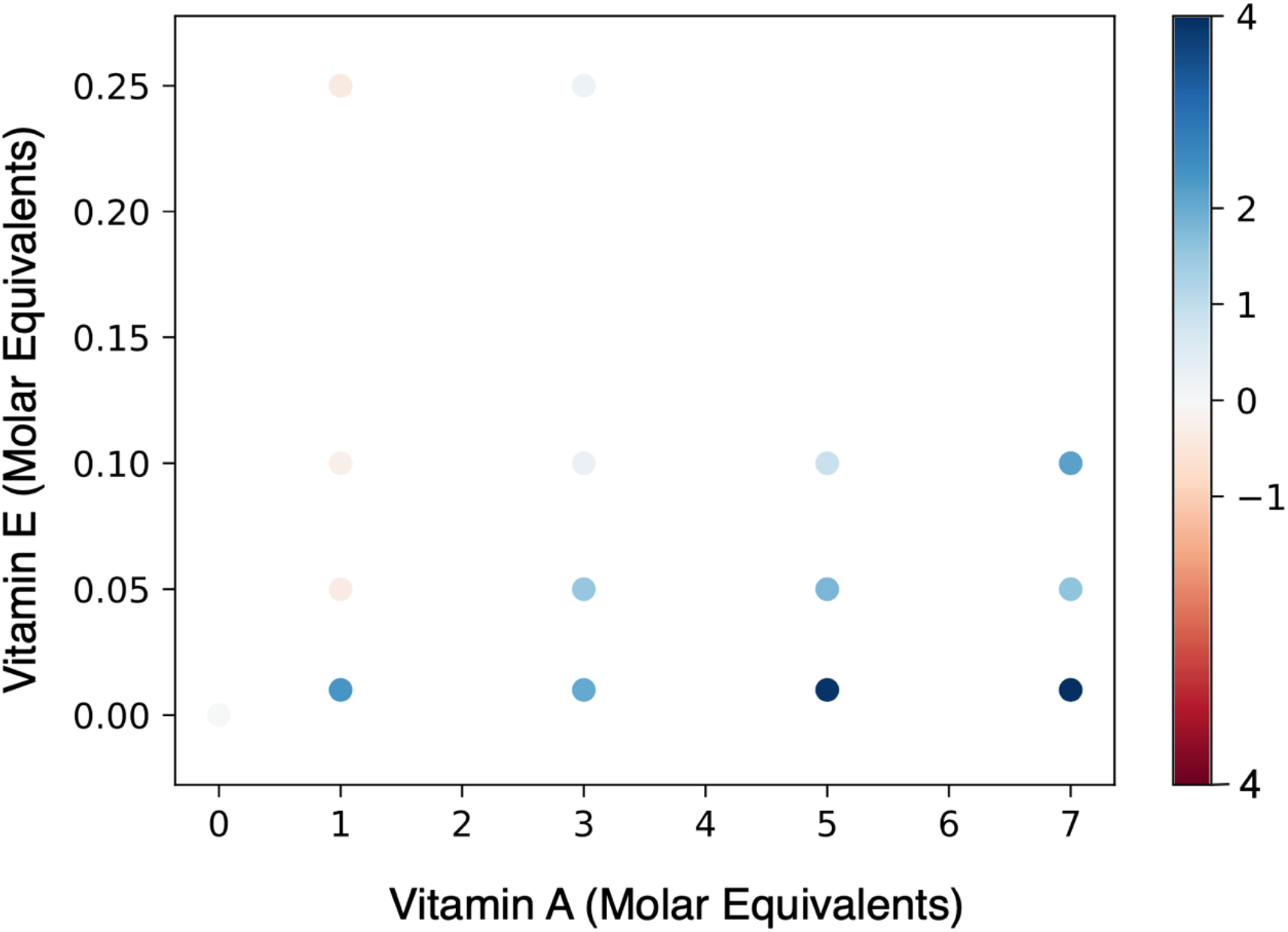
Heat map of the half-time (t_1/2_) of Aβ42 aggregation of different mixtures of vitamin A and vitamin E. The x-axis shows the molar equivalents of vitamin A added to the molar equivalents of vitamin E on the y-axis. The map shows the range of concentrations at which the individual effect of each metabolite is cancelled out by the other, thus having no net effect (zero-sum; equal to all data points representing 0) on Aβ42 aggregation.

To examine these effects independently, we carried out transmission electron microscopy (TEM) on samples deposited on carbon grids in the plateau region (at 4 h) of the aggregation of Aβ42 in the absence and presence of retinoic acid and α-tocopherol (**Figure 6**). We observed a reduction of Aβ42 fibrils in the presence of retinoic acid and an increase of Aβ42 fibrils in the presence of α-tocopherol (**Figure 6**). In the presence of both retinoic acid and α-tocopherol (retinoic acid:α-tocopherol, 7:0.25 ME), we observed Aβ42 fibrils that qualitatively were in the middle of that in the presence of retinoic acid and α-tocopherol respectively.

**Figure 6.**
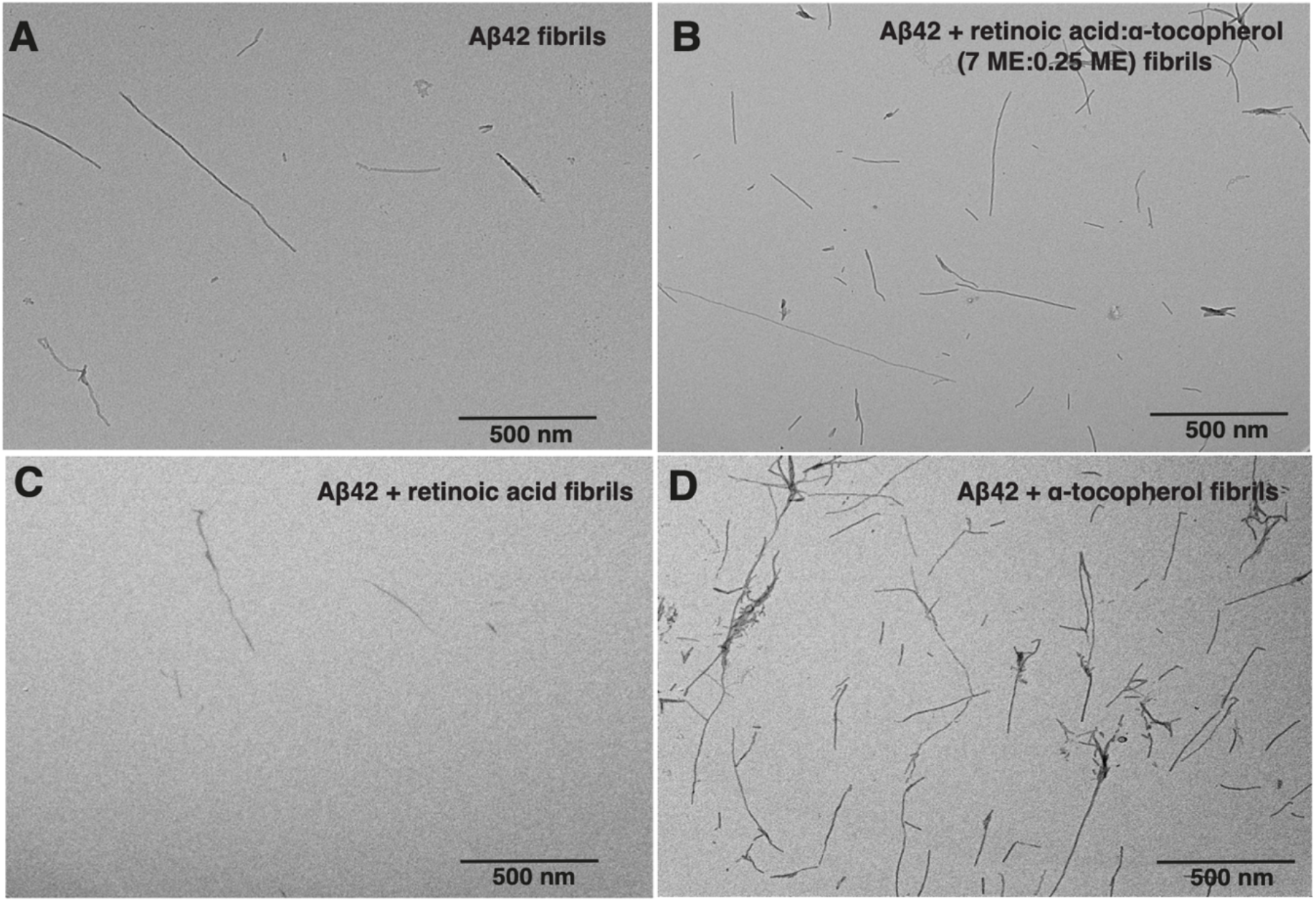
Transmission electron microscopy (TEM) images reveal appearance of a heterogenous population of Aβ42 fibrils after 4 hours incubation at 37 °C in 20mM sodium phosphate buffer in the absence and presence of vitamins/vitamin mixtures. (**A**) Aβ42 alone, (**B**) Aβ42 + Vitamin E (α-tocopherol) and Vitamin A (retinoic acid) mixture (Vitamin A: Vitamin E :: 7 ME: 0.5 ME), (**C**) Aβ42 + retinoic acid (**D**) Aβ42 + α-tocopherol. Images are representative of three observations; three images were taken at different points on the grid. Scale bars represent 500 nm.

**Figure 7.**
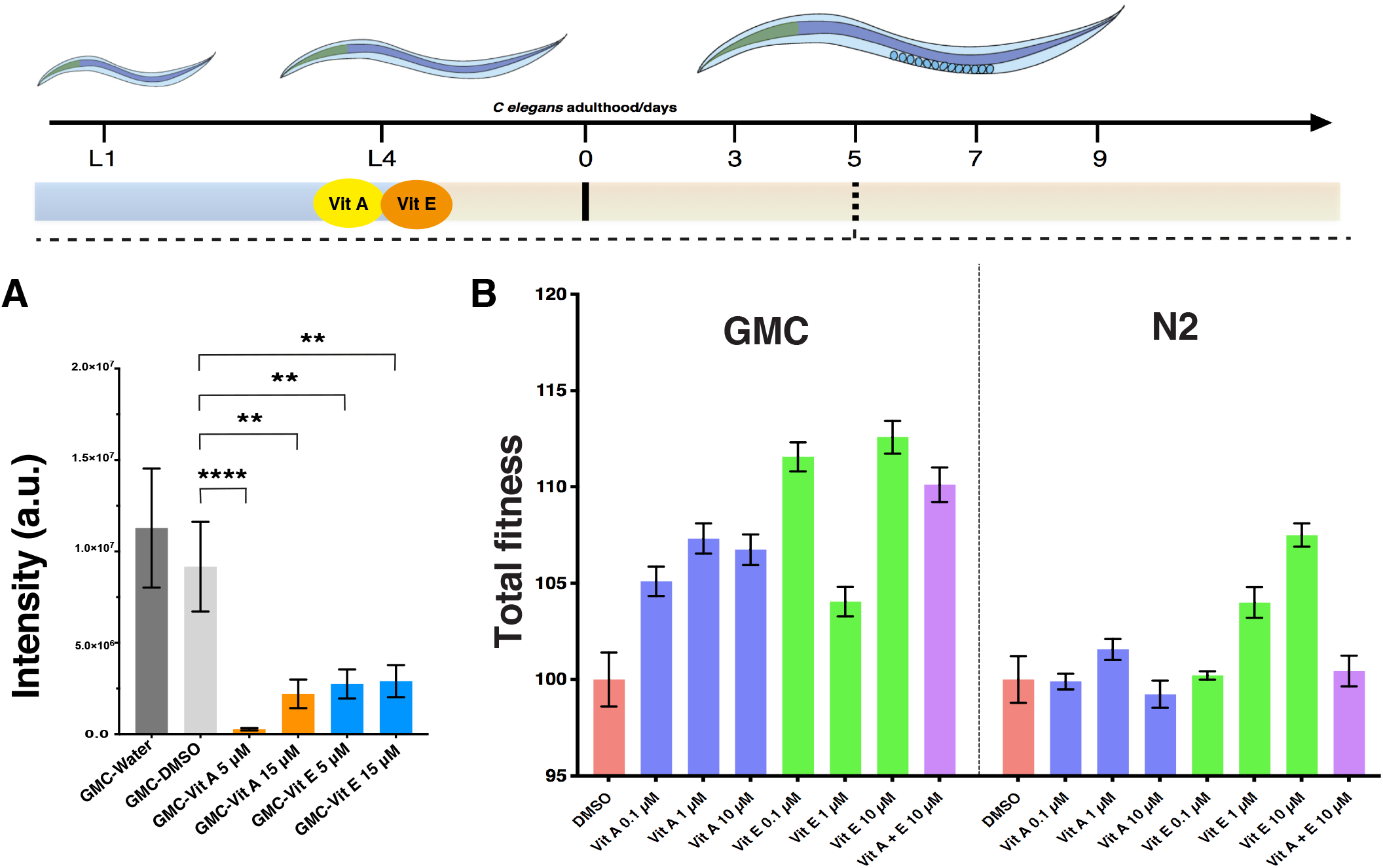
Vitamin A and vitamin E decrease Aβ42 aggregate levels and improve the fitness of a *C. elegans* model of Alzheimer’s disease. We fed with vitamin A and vitamin E L4 worms (GMC strain, represented here as AD for Alzheimer’s disease model) that express Aβ42 in the muscle cells. We observed the effects of the vitamins on the motility of the animals and quantified the aggregate levels by NIAD-4 staining on day 5 of adulthood. **(A)** The intensity of the NIAD-4 staining shows a reduction of aggregates at both 5 and 15 μM of vitamin A and vitamin E, as compared to the control worms treated with water or 1% DMSO. Wild type N2 worms do not exhibit aggregates, hence they are not shown in the panel. **(B)** We treated both wild-type N2 and GMC worms with vitamin A and vitamin E and tracked body bends per minute (BPMs) at day 5. In all cases, we found that the treatment with the two vitamins either increased the fitness of the GMC worms relative to the DMSO control, while the effect was much more limited in the control N2 worms.

These results suggest that taken together the effects of these two vitamins on Aβ42 aggregation can be neutralized when present in combination at a specific range of concentrations. Although here we show only two metabolites exhibiting such an effect, we expect other aggregation promoters and inhibitors to be found that would act similarly in solution. In a proof-of-principle, we thus propose that the complex metabolite-composition of the cellular environment could have a protective role against protein aggregation through the simultaneous presence of aggregation promoters and inhibitors.

### Vitamin A and vitamin E increase the fitness of a *C. elegans* model of Alzheimer’s disease

To study the effects of vitamin A (retinoic acid) and vitamin E (α-tocopherol) on the aggregation of Aβ42 in an *in vivo* system, we used a *C. elegans* model of Alzheimer’s disease. In the GMC101 strain, Aβ42 is expressed in the body wall muscle cells, where it aggregates and results in a severe and fully penetrant, age-progressive paralysis^18^. This gradual aggregation of Aβ42 in the body wall muscle cells causes worm paralysis, which can be quantified by a worm tracking platform^19, 20^. We measured worm motility (body bends per minute) at day 5 of adulthood when the aggregation phenotype is highly visible^15, 19^. Simultaneously, using NIAD-4, which is a dye that specifically binds amyloid fibrils, we measured Aβ42 aggregates in the worm heads^10, 15^ (**Materials and Methods**), and found a decrease in the NIAD-4 stained aggregates for both vitamin A and vitamin E (**Figure 6A**). This reduction of Aβ42 aggregates upon treatment with vitamin A and vitamin E corresponded to an increase in the total fitness of the worms compared to the untreated worms (**Figure 6B**).

## Discussion and Conclusions

Linus Pauling in 1977 proposed that therapies aimed at improving the molecular composition of the brain may restore abnormal functions caused by disease^21^, a concept that was later described as vitamin homeostasis in the brain^22^. Pauling’s suggestion gave rise to the popular megavitamin therapy for disease, which has been the backbone of the global supplements industry for decades despite a lack of clear physiological evidence on health benefits. A major reason for this lack of evidence could be attributed to the absence of fully quantitative methods to mechanistically probe this hypothesis and validate its claims. In this broad context, we sought to investigate the potential roles of vitamins in protein aggregation.

In this work, we have shown that several vitamins can modulate Aβ42 aggregation. Except for vitamin B12, all the vitamins that we identified as positive hits in our screen are fat-soluble. The lipid-like nature of these vitamins could be responsible for their aggregation-modifying behavior, as shown in previous studies on other types of compounds^12, 16, 23, 24^. Further, our results indicate that that the lipid-like nature of these vitamins can lead to the formation of micelles, which in turn may accelerate the aggregation process by enhancing surface-catalysed nucleation (**Figure 3**). The intracellular concentration of this micelle-forming metabolite is thus likely a critical factor deciding whether this pathway of aggregation will proceed or not.

In particular, we have found that vitamin A inhibits the aggregation of Aβ42 in a dose-dependent manner, and that vitamin E accelerates Aβ42 aggregation in its micellar form but inhibits it in a soluble monomeric form. Further, we have described how these two vitamins can have opposite effects on Aβ42 aggregation in a mixture, as we observed a zero-sum effect on Aβ42 aggregation at a range of concentrations. Taken together, we suggest that these vitamins may be added as components of the protein quality control system due to their effects on protein aggregation.

In conclusion, we have shown that vitamin A and vitamin E have opposite effects on Aβ42 aggregation, and that these effects can be cancelled out in a mixture. Taken together, these results suggest that a complex cellular environment may have a protective role through the simultaneous presence of aggregation promoters and inhibitors. If considered as a proof-of-principle, our results illustrate a systematic approach to investigate the possible effects of a dysregulated molecular composition on aggregation-prone proteins. We thus expect that this approach could be extended to other metabolites and proteins implicated in protein misfolding diseases.

## Materials and Methods

### Chemicals

All metabolites were procured from Sigma and were of analytical grade and dissolved in milli-Q grade water (water-soluble) or at a 1% DMSO (lipid/fat-soluble) working concentration. The metabolite stocks were sonicated in a water bath for 2 minutes for effective dissolution of the solutes. All solutions were freshly prepared for each assay and filtered using Anotop 10 0.02 μm filters prior to use in the assays.

#### Expression, purification, and preparation of Aβ42 peptide samples for kinetic experiments

Aβ42 (MDAEFRHDSGYEVHHQKLVFFAEDVGSNKGAIIGLMVGGVVIA) was expressed recombinantly in the *Escherichia coli* BL21 Gold (DE3) strain (Stratagene, CA, USA), and purified as described previously^15^. In the purification procedure, the *E. coli* cells were sonicated, and the inclusion bodies were subsequently dissolved in 8 M urea. A diethyl-aminoethyl cellulose resin was then used to perform ion exchange chromatography, and the protein collected was lyophilized. These fractions were then further purified using a Superdex 75 26/60 column (GE healthcare, IL, USA), and the fractions containing the recombinant protein were combined, frozen and lyophilised again. Solutions of monomeric peptides were prepared by dissolving lyophilized Aβ42 in 6 M GuHCl, and purified in 20 mM sodium phosphate buffer, 200 μM EDTA, pH 8.0 using a Superdex 75 10/300 column (GE Healthcare) at a flow rate of 0.5 ml/min. ThT was added from a 2 mM stock to give a final concentration of 20 μM. All samples were prepared in low-binding Eppendorf tubes and samples were analyzed in a 96-well half area, low-binding, clear-bottom PEG coated plate (Corning 3881). Each sample was then pipetted into multiple wells of a 96-well half-area, low-binding, clear bottom and PEG coating plate (Corning 3881), 80 μL per well, in the absence and the presence of various concentrations of vitamins, to give a final concentration of 1% DMSO (v/v) using an Eppendorf laboratory pipetting robot.

The aggregation of Aβ42, at 2 μM concentration^15, 16^, was initiated by placing the 96-well plate in a plate reader (Fluostar Omega or Fluostar Optima from BMG Labtech, Aylesbury, UK) at 37 °C under quiescent conditions. The ThT fluorescence was monitored in triplicate per sample as measured using bottom-optics with 440 nm excitation and 480 nm emission filters.

#### Transmission electron microscopy (TEM)

Samples were deposited on a 400-mesh, 3mm copper grid carbon support film (EM Resolutions Ltd. Sheffield, UK) and stained with 2% uranyl acetate (w/v). Imaging was carried out on an FEI Tecnai G2 transmission electron microscope (Cambridge Advanced Imaging Centre, University of Cambridge, UK), and the images were acquired using the SIS Megaview II Image Capture system (Olympus, Muenster, Germany). Briefly, the grid was instantaneously washed with chloroform followed by 3x Milli-Q water washes and blotted with a filter paper to remove any debris. 5 μl of sample was deposited on the grid and left for 3 minutes, followed by two instantaneous washes with Milli-Q water. 2.5 μl of uranyl acetate was deposited and left for 2 minutes. This was wicked away using a filter paper to ensure that the patches on the grid are fully stained.

#### *C. elegans* experiments

##### Media

Standard conditions were used for the propagation of *C. elegans*^25^. Animals were synchronized by hypochlorite bleaching (5% hypochlorite solution), hatched overnight in M9 (3 g/l KH_2_PO_4_, 6 g/l Na_2_HPO_4_, 5g/l NaCl, 1 μM MgSO_4_) buffer, and subsequently cultured at 20 °C on nematode growth medium (NGM) (CaCl_2_ 1 mM, MgSO_4_ 1mM, cholesterol 5 μg/ml, 250 μM KH_2_PO_4_ pH 6, Agar 17 g/L, NaCl 3 g/l, casein 7.5 g/l) plates that were seeded with the overnight grown E. coli strain OP50. Saturated cultures of OP50 were grown by inoculating 50 mL of LB medium (tryptone 10 g/l, NaCl 10 g/l, yeast extract 5 g/l) with OP50 and incubating the culture for 16 h at 37 °C. NGM plates were seeded with bacteria by adding 350 μl of saturated OP50 to each plate and leaving the plates at 20 °C for 2-3 days. On day 3 after synchronization, the animals were placed on NGM plates containing 5-fluoro-2’deoxy-uridine (FUdR) (75 μM) to inhibit the growth of offspring and the temperature was raised to 24 °C.

##### Strains

All strains were acquired from the Caenorhabditis Genetics Center in Minnesota, which is supported by NIH P40 OD010440. Two strains were utilised for these experiments. The temperature sensitive human Aβ-expressing strain dvIs100 [unc-54p::A-beta-1-42::unc-54 3’-UTR + mtl-2p::GFP] (GMC101) was used, in which mtl-2p::GFP causes intestinal GFP expression and unc-54p::A-beta-1-42 expresses the human full-length Aβ42 peptide in the muscle cells of the body wall. Raising the temperature above 20 °C at the L4 or adult stage causes paralysis due to Aβ42 aggregation in the body wall muscle. The N2 strain was used for wild-type worms.

##### Vitamin-coated plates

Aliquots of NGM media were autoclaved and poured and seeded with 350 μL OP50 culture that was grown overnight. After incubating for up to 3 days at room temperature, aliquots of vitamin A and vitamin E dissolved in 1% DMSO at different concentrations were added. NGM plates containing FUdR (75 μM) were seeded with 2.2 ml aliquots of vitamins dissolved in water at the appropriate concentration (**Figure 5**). The plates were then placed in a laminar flow hood at room temperature to dry and the worms were transferred to plates coated with the vitamin at larval stage L4. Vitamin A was prepared at room temperature to a stock concentration of 2 mM. Vitamin E stock was prepared at 2 mM. Fresh stocks were made for each experiment.

##### Automated motility assay

At different ages, animals were washed off the plates with M9 buffer and spread over an OP50 unseeded 6 cm plate, after which their movements were recorded at 20 fps using a recently developed microscopic procedure for 90 seconds^19, 20^. Approximately 300 animals were counted at each data point.

##### NIAD-4 staining and imaging

NIAD-4 solution was prepared by dissolution in 100% DMSO at 1 mg/ml. Prior to worm incubation, a 1/1000 dilution in M9 was created. After screening using the Wide-field Nematode Tracking Platform^19, 20^, approximately 300 worms per condition were collected in M9 media and centrifuged at 20 °C at 2000 rpm for two minutes to a pellet. 1 mL of diluted NIAD-4 solution in M9 was then added to the pellet and placed under gentle shaking (80 rpm) for six hours. Worms were then transferred to unseeded NGM plates and incubated at 20 °C for twelve hours. Worms were again washed from the plates with M9 media, spun down, washed with 10 ml M9, and resuspended in 2 ml M9. After gravity sedimentation, 15 μL worm solution was spotted on 4% agarose pads. To anaesthetise the animals, 4 μL of 40 mM NaN3 was added, followed by a glass coverslip. Worms were imaged using a Zeiss Axio Observer A1 fluorescence microscope (Carl Zeiss Microscopy GmbH, Jena, Germany) with a 20x objective and a 49004 ET-CY3/TRITC filter (Chroma Technology Corp, VT, USA). An exposure time of 1,000 ms was employed. The nominal magnification was 40x and images were captured using an Evolve 512 Delta EMCCD camera with high quantum efficiency (Photometrics, Tucson, AZ, USA). Approximately 20 animals were analyzed per condition and statistics were performed using the one-way ANOVA against the untreated group. All statistics herein were preformed using GraphPad Prism. Quantification was preformed using ImageJ (NIH, MD, USA) to determine the grayscale intensity mean in the head of each animal^10^.

---

**Figure S1.**
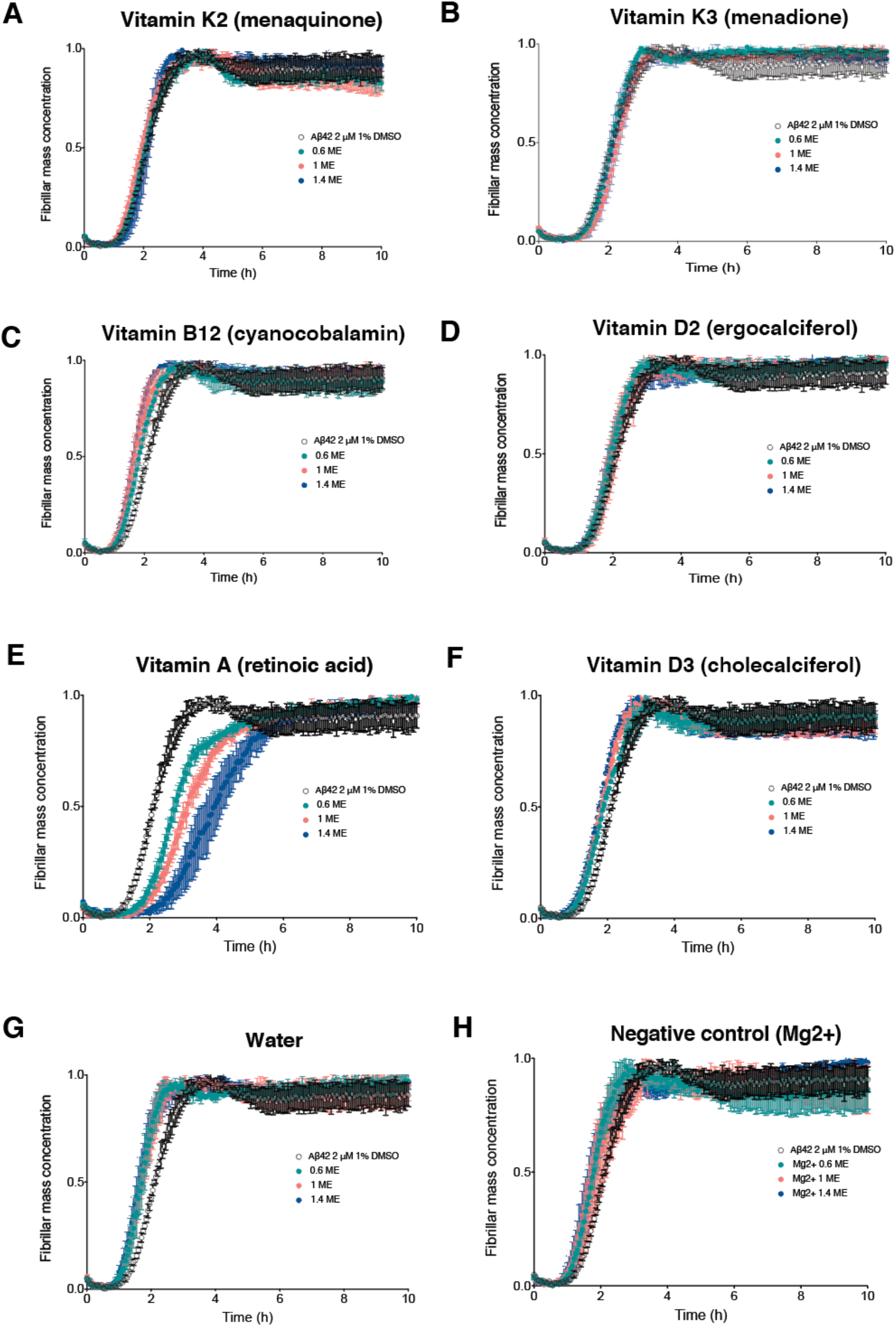
Vitamins D2, D3, K2, K3 and B12 have no effects on Aβ42 aggregation at concentrations below 1.4 ME.

**Figure S2.**
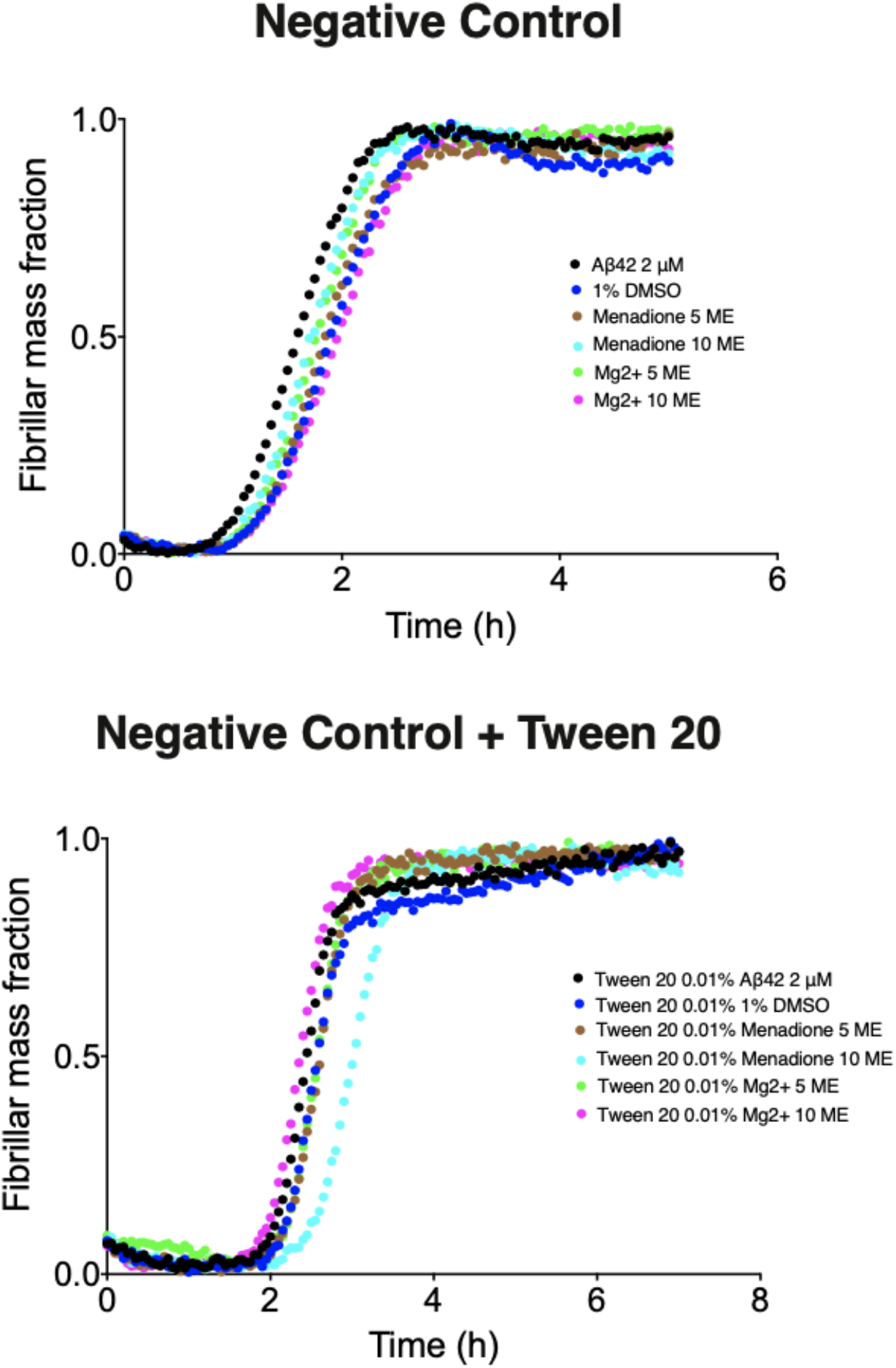
Tween 20 0.01% does not have any effect on Aβ42 aggregation.

## Notes

### Competing Interest Statement

The authors have declared no competing interest.

## References

1. Alzheimer’s Association (2021) Alzheimer’s disease facts and figures. Alzheimer’s & Dementia 7, 327–406.

2. Masters, C. L., Bateman, R., Blennow, K., Rowe, C. C., Sperling, R. A., Cummings, J. L. (2015) Alzheimer’s Disease. Nat. Rev. Dis. Primers 1, 15056.

3. Selkoe, D. J., Hardy, J. (2016) The amyloid hypothesis of Alzheimer’s disease at 25 years. EMBO Mol. Med. 8 (6), 595–608.

4. Knowles, T. P., Vendruscolo, M., Dobson, C. M. (2014) The amyloid state and its association with protein misfolding diseases. Nat. Rev. Mol. Cell Biol. 15 (6), 384–396.

5. Hampel, H., Hardy, J., Blennow, K., Chen, C., Perry, G., Kim, S. H., Villemagne, V. L., Aisen, P., Vendruscolo, M., Iwatsubo, T. (2021) The Amyloid-β Pathway in Alzheimer’s Disease. Molecular Psychiatry, 1–23.

6. Balch, W. E., Morimoto, R. I., Dillin, A., Kelly, J. W. (2008) Adapting proteostasis for disease intervention. Science 319 (5865), 916–919.

7. Hipp, M. S., Park, S.-H., Hartl, F. U. (2014) Proteostasis impairment in protein-misfolding and-aggregation diseases. Trends Cell Biol. 24 (9), 506–514.

8. Lindquist, S. L., Kelly, J. W. (2011) Chemical and biological approaches for adapting proteostasis to ameliorate protein misfolding and aggregation diseases–progress and prognosis. Cold Spring Harbor Perspect. Biol. 3 (12), a004507.

9. Hipkiss, A. R., Cartwright, S. P., Bromley, C., Gross, S. R., Bill, R. M. (2013) Carnosine: can understanding its actions on energy metabolism and protein homeostasis inform its therapeutic potential? Chem. Cent. J. 7 (1), 38.

10. Joshi, P., Perni, M., Limbocker, R., Mannini, B., Casford, S., Chia, S., Habchi, J., Labbadia, J., Dobson, C. M., Vendruscolo, M. (2021) Two human metabolites rescue a C. elegans model of Alzheimer’s disease via a cytosolic unfolded protein response. Comm. Biol. 4 (1), 1–14.

11. Choi, J., Yin, T., Shinozaki, K., Lampe, J. W., Stevens, J. F., Becker, L. B., Kim, J. (2018) Comprehensive analysis of phospholipids in the brain, heart, kidney, and liver: brain phospholipids are least enriched with polyunsaturated fatty acids. Mol. Cell. Biochem. 442 (1-2), 187–201.

12. Sanguanini, M., Baumann, K. N., Preet, S., Chia, S., Habchi, J., Knowles, T. P., Vendruscolo, M. (2020) Complexity in lipid membrane composition induces resilience to Aβ42 aggregation. ACS Chem. Neurosci. 11 (9), 1347–1352.

13. Fenech, M. (2017) Vitamins associated with brain aging, mild cognitive impairment, and Alzheimer disease: Biomarkers, epidemiological and experimental evidence, plausible mechanisms, and knowledge gaps. Adv. Nutr. 8 (6), 958–970.

14. Cohen, S. I., Vendruscolo, M., Dobson, C. M., Knowles, T. P. (2012) From macroscopic measurements to microscopic mechanisms of protein aggregation. J. Mol. Biol. 421 (2-3), 160–171.

15. Habchi, J., Arosio, P., Perni, M., Costa, A. R., Yagi-Utsumi, M., Joshi, P., Chia, S., Cohen, S. I., Müller, M. B., Linse, S. (2016) An anticancer drug suppresses the primary nucleation reaction that initiates the production of the toxic Aβ42 aggregates linked with Alzheimer’s disease. Sci. Adv. 2 (2), e1501244.

16. Habchi, J., Chia, S., Galvagnion, C., Michaels, T. C., Bellaiche, M. M., Ruggeri, F. S., Sanguanini, M., Idini, I., Kumita, J. R., Sparr, E. (2018) Cholesterol catalyses Aβ42 aggregation through a heterogeneous nucleation pathway in the presence of lipid membranes. Nat. Chem. 10 (6), 673–683.

17. Meisl, G., Kirkegaard, J. B., Arosio, P., Michaels, T. C., Vendruscolo, M., Dobson, C. M., Linse, S., Knowles, T. P. (2016) Molecular mechanisms of protein aggregation from global fitting of kinetic models. Nat. Protoc. 11 (2), 252–272.

18. McColl, G., Roberts, B. R., Pukala, T. L., Kenche, V. B., Roberts, C. M., Link, C. D., Ryan, T. M., Masters, C. L., Barnham, K. J., Bush, A. I. (2012) Utility of an improved model of amyloid-beta (Aβ 1-42) toxicity in Caenorhabditis elegans for drug screening for Alzheimer’s disease. Mol. Neurodegener. 7 (1), 1–9.

19. Perni, M., Casford, S., Aprile, F. A., Nollen, E. A., Knowles, T. P., Vendruscolo, M., Dobson, C. M. (2018) Automated behavioral analysis of large C. elegans populations using a wide field-of-view tracking platform. JoVE (141), e58643.

20. Perni, M., Challa, P. K., Kirkegaard, J. B., Limbocker, R., Koopman, M., Hardenberg, M. C., Sormanni, P., Müller, T., Saar, K. L., Roode, L. W. (2018) Massively parallel C. elegans tracking provides multi-dimensional fingerprints for phenotypic discovery. J. Neurosci. Methods 306, 57–67.

21. Pauling, L. (1974) On the orthomolecular environment of the mind: orthomolecular theory. Am. J. Psychiatry 131 (11), 1251–1257.

22. Spector, R. (1977) Vitamin homeostasis in the central nervous system. N. Engl. J. Med. 296 (24), 1393–1398.

23. Hellstrand, E., Sparr, E., Linse, S. (2010) Retardation of Aβ fibril formation by phospholipid vesicles depends on membrane phase behavior. Bioph. J. 98 (10), 2206–2214.

24. Lau, T.-L., Gehman, J. D., Wade, J. D., Masters, C. L., Barnham, K. J., Separovic, F. (2007) Cholesterol and Clioquinol modulation of Aβ (1–42) interaction with phospholipid bilayers and metals. Biochim. Biophys. Acta, Biomembr. 1768 (12), 3135–3144.

25. Brenner, S. (1974) The genetics of Caenorhabditis elegans. Genetics 77 (1), 71–94.

